# Celiac dysbiosis does not transcend geographic boundaries

**DOI:** 10.1101/2022.10.02.510531

**Authors:** John J Colgan, Michael B Burns

**Affiliations:** Loyola University Chicago

## Abstract

Celiac disease is an autoimmune disorder of the small intestine in which gluten, an energy-storage protein found in wheat and other cereals, elicits an immune response that leads to villous atrophy. Despite a strong genetic component, celiac disease arises sporadically and at any age, leading us to hypothesize that changes in the microbiome influence celiac disease development and/or progression. Here, we pooled and computationally analyzed 16S data from 3 prior international studies that examined celiac disease and the microbiome. For our analysis, we combined the dada2 and PICRUSt 2 pipelines and a variety of data transformations that control for batch effects to determine whether any taxonomic or metabolic features were consistently associated with the celiac microbiome across the globe. Our results showed the celiac microbiome displays dysbiosis without a discernable pattern, which suggests perturbations in the celiac microbiome are a result of the disease rather than a cause. Data from PICRUSt 2 supported this conclusion and revealed connections between celiac disease and the metabolome that are supported by previous research examining dysbiotic microbiomes.

**IMPORTANCE:** Celiac disease is an autoimmune disorder that affects roughly 2% of the world’s population. Although the ultimate cause of celiac disease is unknown, many researchers hypothesize that changes to the intestinal microbiome play a key role in disease progression. If this is the case, it may be possible to design therapies that manipulate the microbiome to suppress celiac disease. Here, we analyzed pooled data from 3 different studies from across the globe that examined celiac disease and the microbiome to ascertain whether there exists a unique celiac microbiome that transcends geographic boundaries.

## INTRODUCTION

Celiac Disease (CD) is an inflammatory bowel disease (IBD) of the small intestine in which the protein gluten from wheat and barley causes an inflammatory response that degrades the lining of the small intestine, specifically the villi (1). If CD is left untreated, patients acquire a host of health problems, including malnutrition, osteoporosis and, in rare cases, cancer (1). CD is estimated to affect up to 2% of the world’s population, with Western countries having the highest incidence. However, many cases of celiac disease go undiagnosed (1). CD can present at almost any age with intestinal and non-intestinal symptoms (1). The only existing treatment for CD is following a gluten-free diet (GFD), which excludes gluten and thereby prevents the inflammatory response (1).

All CD patients have one or both of the HLA DQ2 and DQ8 haplotypes. However, concordance between CD and these haplotypes is not 100% (2) suggesting environmental factors play a role in CD progression. Recent research found CD subjects have a dysbiotic gut microbiome, but no clear pattern for the celiac microbiome was identified (1). Although several studies found evidence that the celiac microbiome harbors an excess of bacterial taxa associated with inflammation, no specific bacterium or microbial community configuration was directly implicated (1). The characteristic inflammatory response seen in celiac disease could be caused by the metabolic activity of the implicated microbes, whereby byproducts of microbial metabolism modulate the immune system. This type of effect was demonstrated for the short chain fatty acid butyrate, which is produced by bacterial digestion of fiber. In mice, butyrate was shown to ameliorate symptoms of rheumatoid arthritis, another autoimmune disorder, thus demonstrating the ability of symbiotes to modulate the immune system (3).

In our previous study, we used newer computational tools to reanalyze data from older CD studies to determine whether these tools would provide additional insights (4). Here, we analyzed pooled data from 3 previous international studies to look for associations between the microbiome and CD and determine whether any taxa or metabolic pathways are associated with CD. Our analysis was conducted using the dada2 pipeline (5) to generate taxonomic classifications for each read. These data were then passed off to Clustal-o (6) and Fast-Tree (7) to create a phylogenetic tree and PICRUSt 2 (8-11) to obtain functional analysis of the microbes. These data were then analyzed using microbiome analyst to obtain alpha- and beta-diversity metrics and identify differentially abundant taxa and metabolic pathways (12).

We included data from Bonder et al. (13), which examined stool samples from 21 participants, from Garcia-Mazcorro et al. (14) which defined the microbiomes of 12 celiac patients, 12 non-celiac, gluten-sensitive patients, and 12 controls on GD and GFD via paired stool and duodenum samples, and from Bodkhe et al. (15) which studied 23 untreated celiac patients, 15 first-degree relatives without celiac disease and 24 controls with hepatitis B or functional dyspepsia via paired stool and duodenal biopsies. To supplement these data, 19 healthy stool samples from Chaudhari *et al*. (16) and 17 from Dubey *et al*. (17) were used in the analysis of the stool samples from Bodkhe *et al*. From across these studies we were able to incorporate 166 participants, with 31 celiac patients, 12 non-celiac gluten sensitive patients, 15 first-degree relatives, 24 patients with functional dyspepsia or hepatitis B, and 84 healthy controls, making this work one of the largest meta-analysis examining CD across both the duodenal and stool microbiomes.

## MATERIALS AND METHODS

Studies examining the celiac microbiome were identified using the search queries “celiac microbiome”, “celiac disease and the microbiome”, “celiac disease and gut microbiota”, and “celiac disease and gut-microbiome”. The selected studies reported microbiome sequencing data for the v4 variable region of 16S ribosomal subunit (rRNA). Sequences were prepared for analysis with dada2 (5) by removing adapter sequences with Cutadapt (18). Adapter sequences were provided in the Materials and Methods sections of the respective parent study with taxonomy assigned using the Silva nr99 v138 training set (19). The amplicon sequence variant (ASV) tables from each study were then split into biopsy and stool sample sets that were each analyzed separately. To create a phylogenetic tree, Clustal-o (6) was used locally, with the alignment being passed off to FastTree (7).

The data generated by dada2 were prepared for PICRUSt2 by creating a .fasta file of the ASV sequences and .biom table utilizing the BIOM R package (20). Data were then passed to PICRUSt2 (8 - 11) and analyzed using the default parameters. The resulting data were then passed to microbiomeAnalyst (21, 22). Filtering in microbiomeAnalyst was done in accordance with each respective parent study’s methods without any transformation or refraction. Default parameters were used for weighted unifrac, unweighted unifrac, Shannon diversity index, Simpson diversity index, Chao1 diversity index, RNA seq (EdgeR, 24 -26) and metagenome seq (27-28).

For LEFSe analysis (29), features with a P-value (unadjusted) less than 0.1 and LDA score with an absolute value of 2 or more were identified as significant. Random forest was also applied, which failed to identify any significant taxa accurately. Before filtering, ASVs without taxonomic assignment below Kingdom level were excluded, which removed 23,336 sequences. The data were filtered to remove ASVs with counts less than 4 or prevalence in less than 10% of samples. Features with a variance of less than 10% based on the interquartile range were also discarded. This removed 5065 ASVs and left 6684 ASVs for the remaining analysis. 5 samples with a library size of less than 3000 ASVs were also excluded. The data were then analyzed according to the procedure described in Gibbons *et al*. (23)., with filtering and total sum scaling being applied. Analysis of pathway data was done using Bray-curtis, RNA seq, metagenomeSeq, and LEFSe, all of which were run using default parameters.

## RESULTS

### Pooled, filtered, and scaled analysis of duodenum microbiome samples

Unassigned taxa were removed prior to filtering, leaving 6684 ASVs. Filtering removed 4681 low abundance features and 201 low variance features leaving 1802 features for analysis. Additionally, total sum scaling was applied.

### Community structure analysis

When looking at a factor of disease, Chao1 analysis showed that non-inflammatory bowel disease samples (NIBD) had the highest alpha-diversity and healthy samples had less diversity, with CD samples falling in between (p = 0.27608 ANOVA). Shannon analysis showed that CD and healthy samples had similar alpha-diversities, with NIBD samples having the highest (p = 0.48161 ANOVA). Simpson diversity index showed that CD had the lowest average alpha-diversity (p =0.19143 ANOVA, Figure 1A). Chao1 and Shannon indices showed that CD and non-CD samples had similar alpha-diversities (p = 0.83982, p = 0.91247 ANOVA) with Simpson showing CD samples with a lowered average alpha-diversity (p = 0.30967 ANOVA, Figure 1B). None of these results were significant. All metrics showed that Indian samples had greater alpha-diversity, though this was only significant for Chao1 analysis (p = 0.027265 ANOVA, Figure 1C). Clustering was only achieved based on geographic region rather than disease or CD status for both weighted and unweighted unifrac (p< 0.001 PERMANOVA). No clustering was apparent as a factor of disease or CD status (unweighted unifrac disease p< 0.001, weighted unifrac disease p < 0.001, unweighted unifrac CD status p < 0.133, weighted unifrac p < 0.124 PERMANVOA)

**Figure 1.**
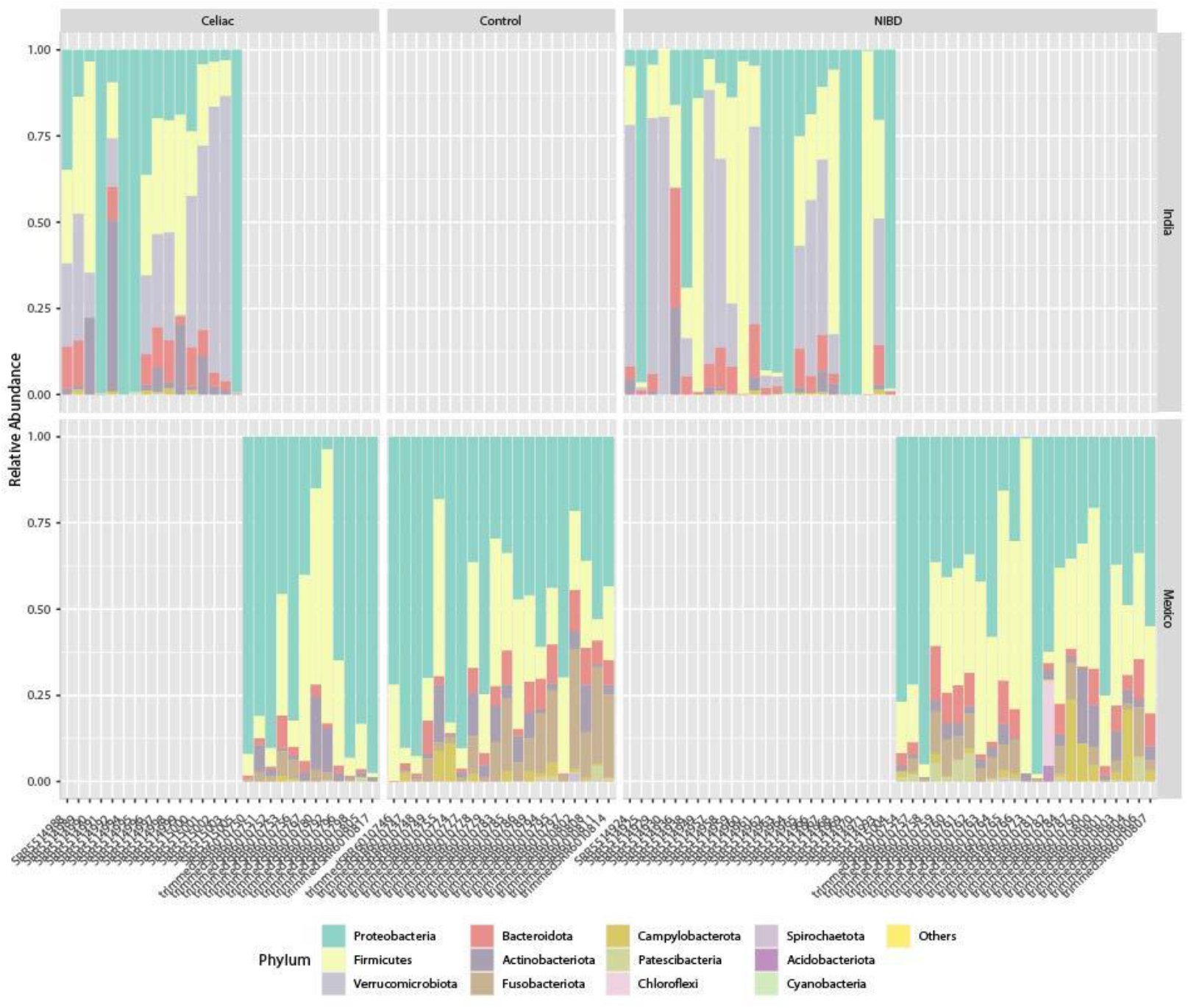
pooled filtered and scaled duodenal abundance plots: Taxonomic distribution of filtered and scaled duodenal biopsies at the phylum level. Shown are percent abundance stacked area barplots of duodenum biopsies fromIndian CD and NIBD patients and Mexican CD, healthy, and NIBD patients at the phylum level. Differences in community structure were noted as both a factor of disease and region of isolation.

Stacked area bar plots of percentage abundance showed differences in community structure in accordance with disease and region. Indian samples contained large proportions of *Verrucomicrobitoa*. Indian CD samples were characterized by an abundance of *Bacteriodata* with Mexican CD samples having the opposite trend. Mexican samples contained *Fusobacteriota*, which was largely absent from Indian samples, with Mexican CD samples having lowered abundance compared to NIBD and controls (Figure 2).

**Figure 2.**
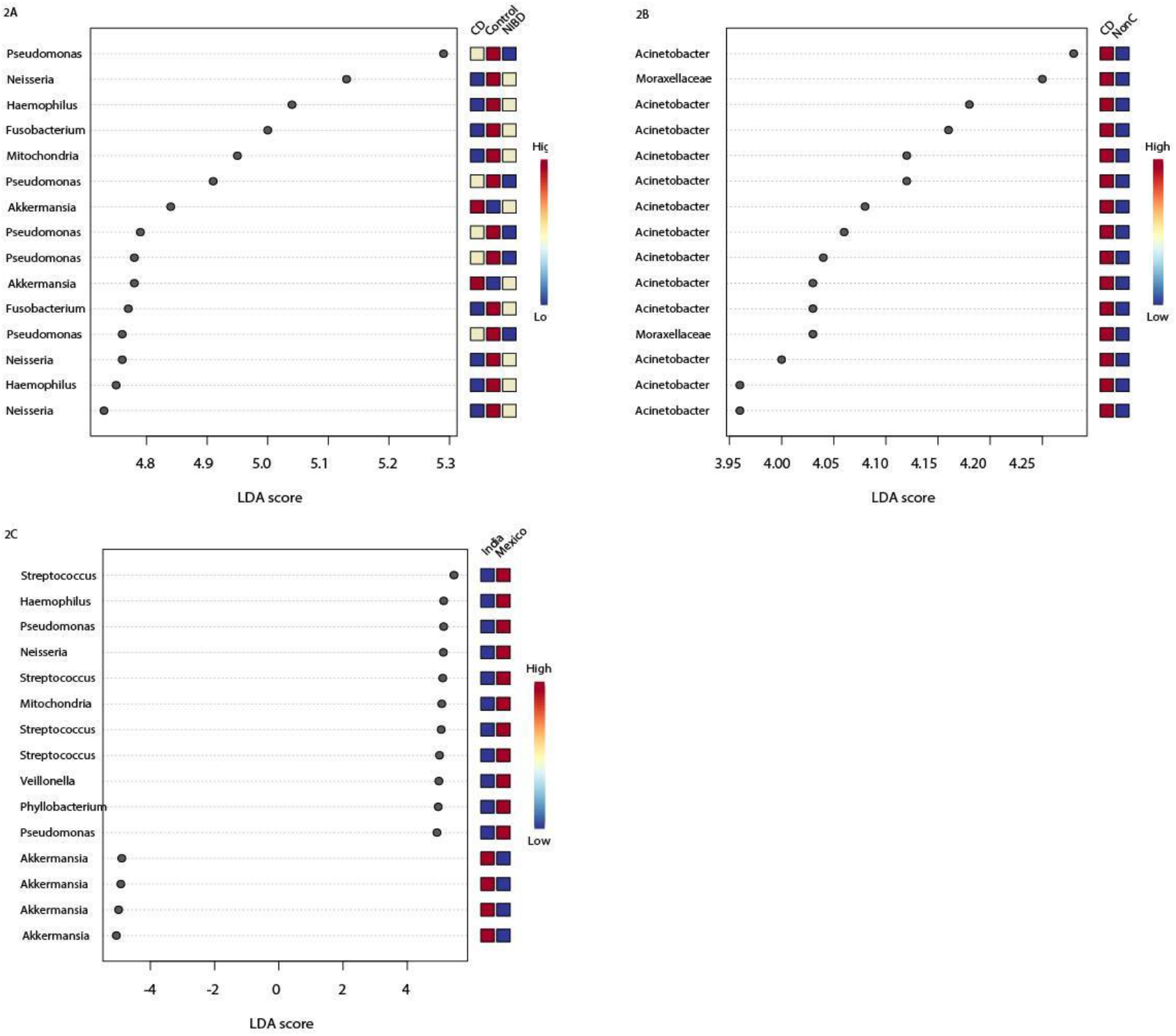
pooled filtered and scaled duodenal LEFSe results: Duodenum filtered and scaled differentially abundant taxa. A) shows differentially abundant taxa identified by LEFSe as a function of disease. B) shows differentially abundant taxa as a function of CD versus non-CD. C) shows differentially abundant taxa as a function of region. Red indicates a higher relative abundance, while beige indicates a similar abundance to red. Blue indicates a lower abundance relative to the group with the highest relative abundance. All taxa had LDA scores > 2.0 and P / FDR lower than 0.1.

### LEFSe

LEFSe identified 107 differentially abundant taxa. Four *Pseudomonas* ASVs were elevated in CD and controls but lowered in NIBD. Three *Neisseria* ASVs were lowered in CD and elevated in controls and NIBD. Two *Haemophilus* ASVs were elevated in controls and for NIBD 2 *Fusobacterium* ASVs were lowered in abundance for CD and 2 *Akkermansia* ASVs were elevated in CD and NIBD, (Figure 3A). 25 ASVs were identified as differentially abundant when looking at CD versus non-CD samples. Of the 15 LDA highest score taxa, 13 ASVs were identified as *Acinetobacter*, and two as belonging to the family *Moraxellaceae* (Figure BB). When examining region of isolation, 500 significant ASVs were identified, with overlap of previously identified genre occurring with *Haemophilus* and *Neisseria*, both of which were associated with the Mexican cohort microbiome, and *Akkermansia*, which was associated with the Indian cohort microbiome (Figure 3C). All ASVs had LDA scores of 2.0 or greater and FDR-adjusted p-values less than or equal to 0.1.

**Figure 3.**
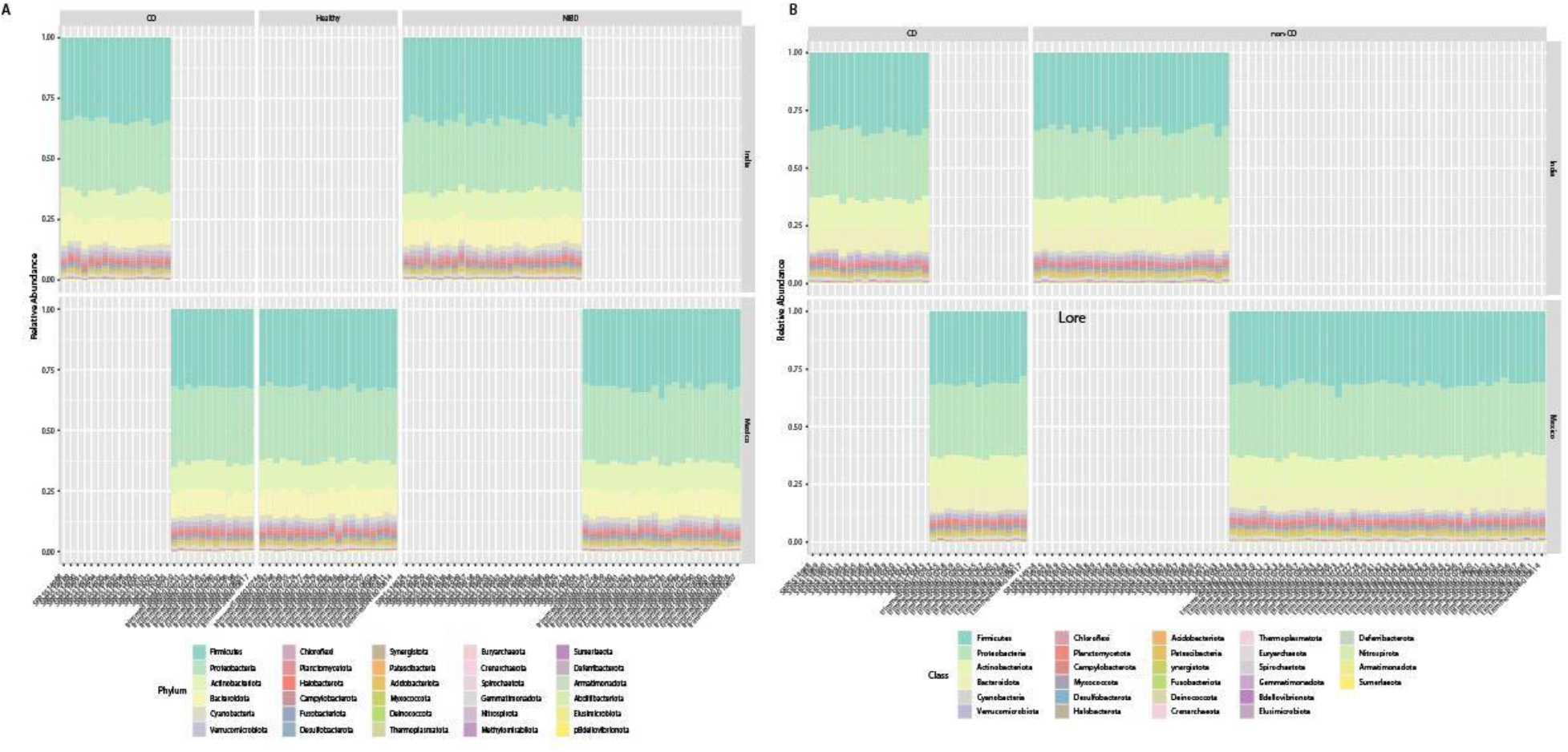
Normalized abundance plots. A) Control normalized abundance plots of Indian and Mexican duodenal biopsies. B) Non-celiac normalized abundance plots of Indian and Mexican biopsies.

### Pooled normalized duodenum samples

The normalization procedure from Gibbons et al. left 799 ASVs out of a total of 6684. After filtering 719, taxa remained with 80 low variance features removed.

### Community structure analysis

Both the normalized control and non-CD duodenum samples had significantly lowered alpha-diversity for diseased patients (Shannon control normalized p = 7.8156*10^−8^, Simpson control normalized p = 7.8156*10^−5^, Shannon non-CD normalized p = 3.5834*10^−6^, Simpson non-CD normalized p = 0.0001319, Shannon non-CD normalized p = 0.034427 ANOVA, Figure 4A, B). No clustering was achieved using weighted unifrac for either normalization group (p > 0.05 PERMANOVA). Interestingly, the results from unweighted unifrac showed many samples falling on the same point in the plane when plotting the first two principal coordinates. This pattern was observed for both the normalized data sets and most dramatic for normalized control samples; where all healthy samples collapsed as a single point. This resulted in disease samples being scattered around the controls. To determine whether this represented clustering, other analysis methods were used (Bray-Curtis, Shannon-Jenson, Jaccard). None of these methods generated clusters (p> 0.05 PERMANOVA, Figure 4 A, C). No differences were noted in the stacked percent abundance bar plots for either normalization set (Figure 5A, B).

**Figure 4.**
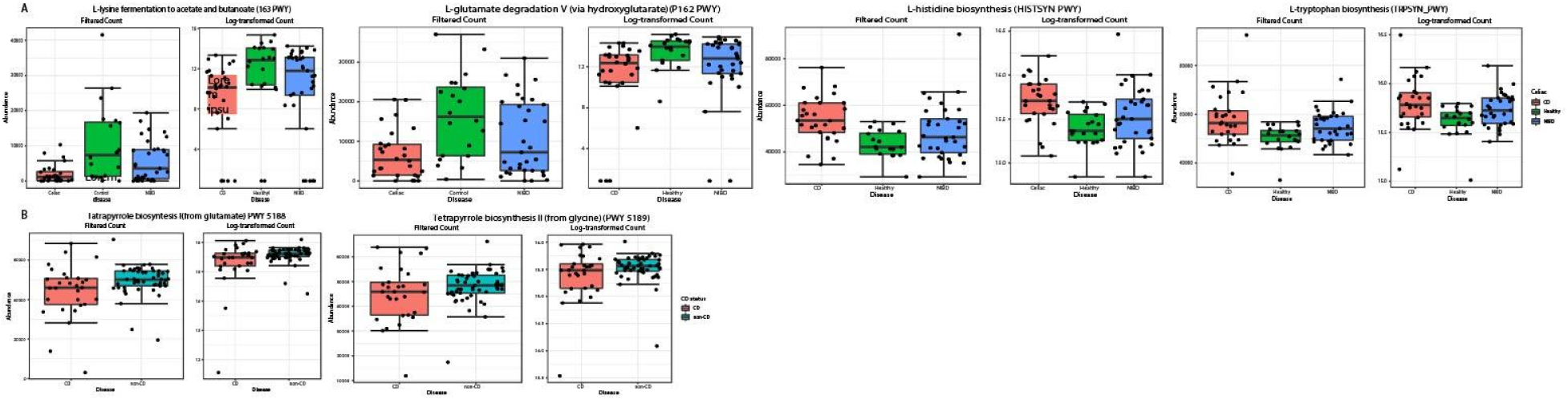
pooled differentially abundant duodenal metabolic pathways: Figure A differentially abundant pathways as a factor of disease with CD shown in red, NIBD blue and controls green. All pathways were identified as significant across LESFSe (LDA > 2.0) as well as metagenome and RNA seq (P > .1). Figure 4B differentially abundant pathways as a factor of CD status with CD in red and non-CD in blue. All pathways were identified as significant using LEFSe (LDA >2.0) RNA and metagenome seq (P >0.1).

**Figure 5.**
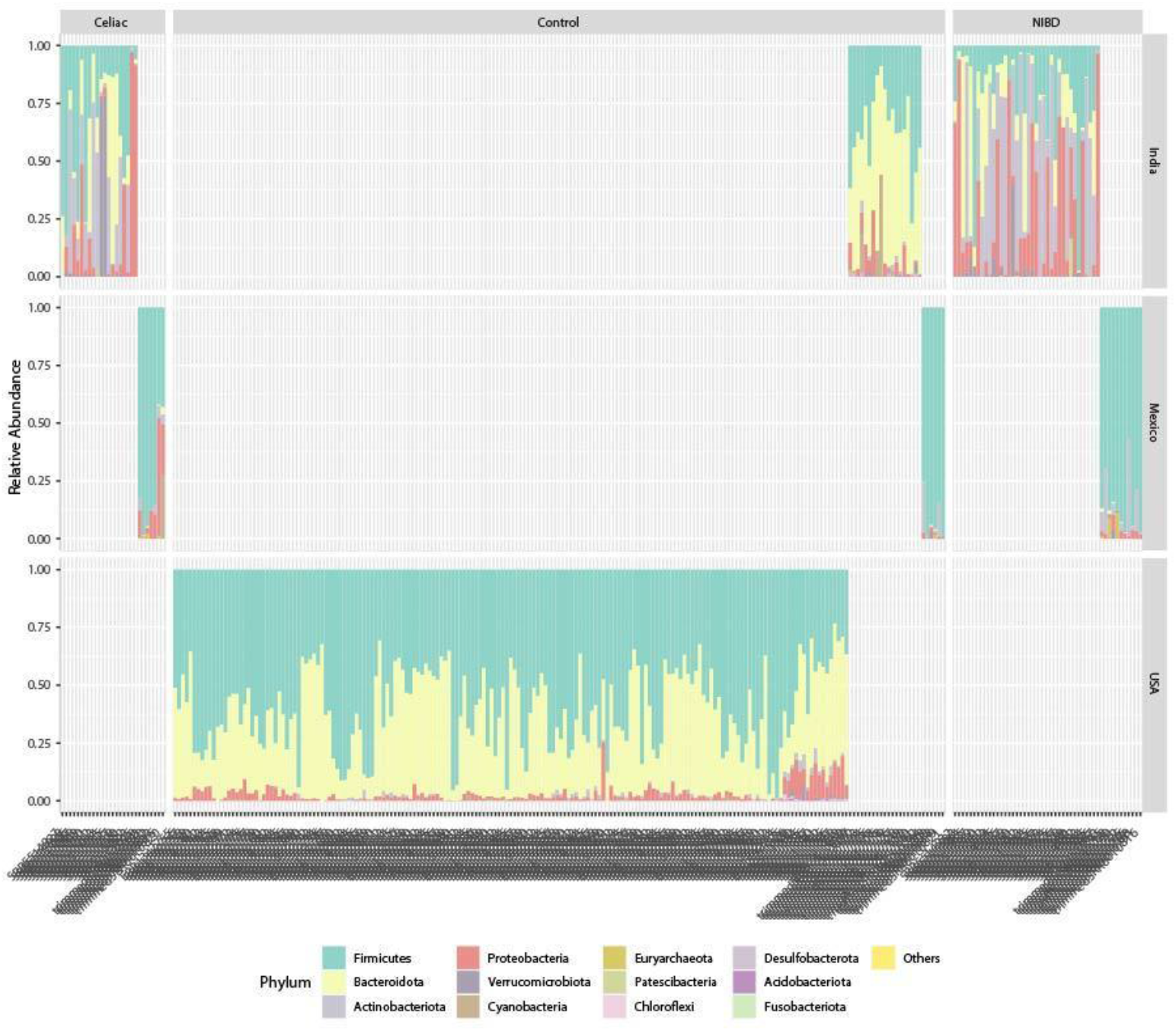
Filtered and scaled fecal abundance plots: Stacked percent abundance bar plots of filtered and scaled data showing community structure at the phylum level. Differences were noted due to both region and disease state.

### LEFSe

No significant taxa were identified using LEFSe for either normalization group.

### Duodenum PICRUSt 2 Pathways

PICRUSt 2 data were analyzed using microbiomeAnalyst’s default parameters. This dataset was much smaller and far less noisy than the ASV data generated by dada2, which made normalization unnecessary. No samples were excluded due to low library size. 12 low abundance features and 41 low variance features were removed using default settings, leaving 368 features.

### Clustering

No clusters were generated based on disease, CD status or geography (Bray-Curtis disease p < 0.034, Bray-Curtis CD status p < 0.097, Bray-Curtis region p < 0.001 PERMANOVA).

### Differentially abundant pathways

Pathway analysis identified 16 shared pathways between RNA-seq (EdgeR), metagenomeSeq, and LEFSe. Pathways that were lowered in CD samples were for norspermidine biosynthesis (PWY 6562), L-lysine fermentation to acetate and butanoate (P163 PWY), methylaspartate cycle (PWY 6728), L-glutamate degradation V (P162 PWY), UDP-2, 3-acetamido-2, 3-dideoxy-a-D-mannuronate biosynthesis (PWY 7090), and glycogen degradation III (PWY 5767) lowered. Pathways for L-ornithine biosynthesis I (GLUTORN), L-histidine biosynthesis (HISTSYN), methanogenesis from acetate (METH-ACETATE PWY), dTDP-N-acetylhomosamine biosynthesis (PWY 7315), L-tryptophan biosynthesis (TRPSYN), and guanosine deoxyribonucleotides de novo biosynthesis II (PWY 7222) were elevated in CD/ NIBD samples and lowered in controls (RNA seq/ metagenomeSeq p < 0.1, LEFSe LDA > 2.0, Figure 6).

**Figure 6.**
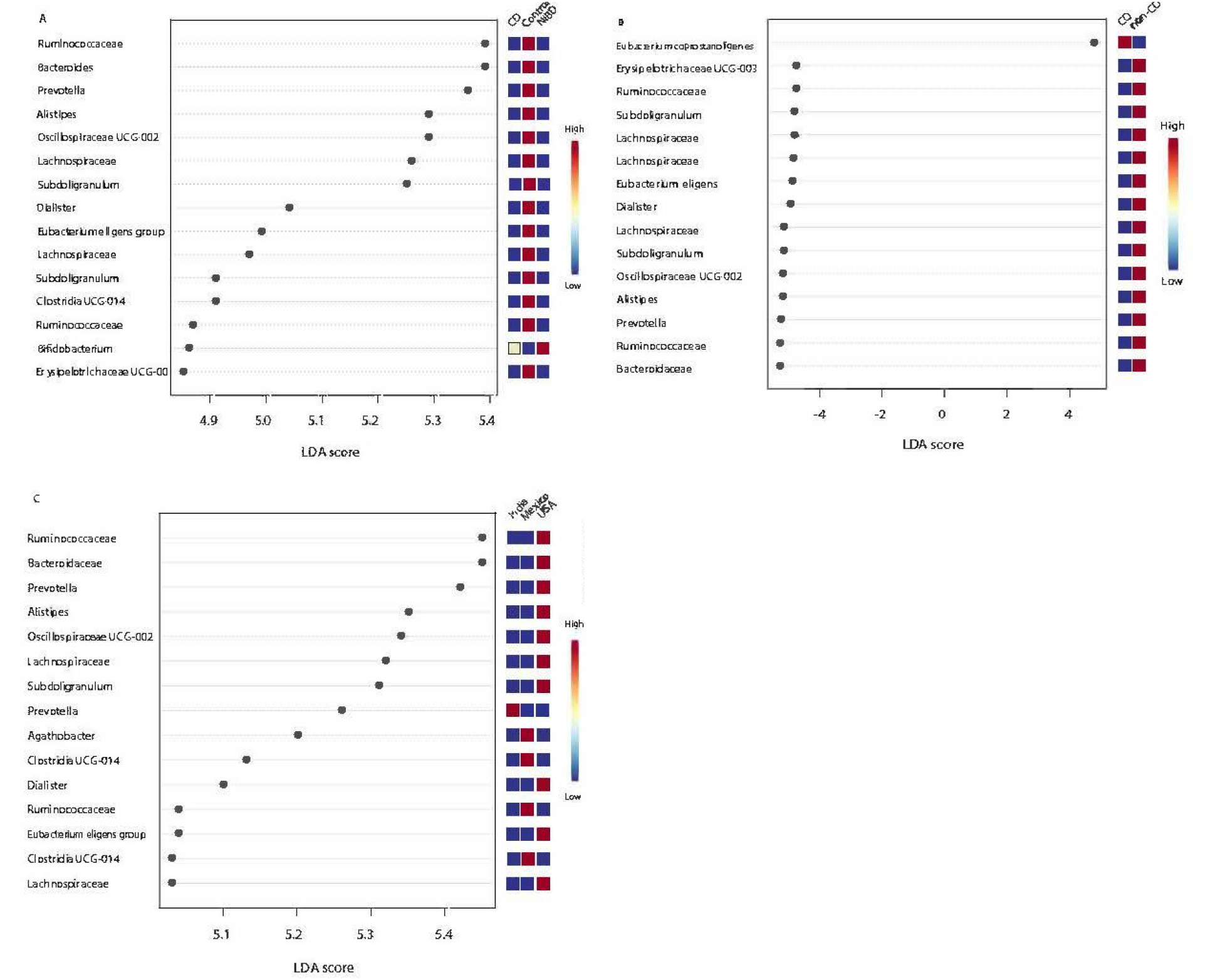
LEFSe differentially abundant filtered and scaled stool taxa. A) show.; differentially abundant taxa as a factor of disease.Bl shows differentially abundant taxa as a function of CD status. C) shows differentially abundant taxa as a function of sample region. Red indicates an elevated relative abundance, beige a similar abundance to red, and blue lowered relative abundance. All features had an LDA score greater than 2.0, P and FDR of less than 0.1.

When comparing CD and non-CD samples, 9 features (UMP biosynthesis (PWY 5686), L-glutamate degradation V (via hydroxyglutarate)(P162 PWY, guanosine deoxyribonucleotides de novo biosynthesis (PWY 6125), and super pathway of purine nucleotides de novo biosynthesis I (PWY 841)) were higher in CD, while pathways of arginine and polyamine biosynthesis (ARG+polyamineSYN), tetrapyrrole biosynthesis I from glutamate (PWY 5188), L-lysine fermentation to acetate and butanoate (P163 PWY), reductive acetyl coenzyme A pathways I (homoacetogenic bacteria)(CODH PWY), and tetrapyrrole biosynthesis from glycine (PWY 5189) were elevated in non-CD. All pathways had LDA scores with an absolute value of 2.5 or greater and RNA(EdgeR)/metagenomeSeq p < 0.1 (Figure 6).

### Pooled, filtered, and scaled analysis of feces microbiome samples

Unassigned taxa were removed prior to filtering, leaving 6684 ASVs. Filtering removed 4689 low-abundance features and 200 low-variance features, leaving 1795 ASVs, with total sum scaling applied.

### Community structure analysis

Chao1, Shannon, and Simpson analysis showed NIBD samples had an elevated alpha-diversity compared to both CD and healthy samples (p = 7.1026*10^−28^, p = 8.0086*10^−23^, p = 3.3789*10^−13^, ANOVA, Figure 7A). CD samples had higher alpha-diversity compared to non-CD across metrics (p =0.0090813, p = 0.0015761, p = 3.357*10^−7^ ANOVA, Figure 7B). Similarly, Indian samples were enriched across metrics (p = 3.1866*10^−21^p = 1.31478*10^−11^, p = 8.9231*10^−7^ ANOVA, Figure 7C). Only weighted unifrac was able to produce significant clusters based on geography, indicating that low-abundance taxa were mainly responsible for the separation seen (p< 0.001, Figure 7D-F). Stacked area plots of percent abundance showed that Indian samples had the most varied community structure with healthy samples having larger proportions of *Bacteriodota*, NIBD samples having a larger proportion of *Actinobacteria* and CD samples having *Proteobacteria*. Mexican CD patients also had elevated *Proteobacteria* compared to other Mexican study groups, though not as enriched compared to the Indian samples. Healthy samples from the United States were homogenous in community structure and dominated both *Bacteroidetes* and *Fimicutes* with small populations of *Proteobacteria* (Figure 8).

**Figure 7.**
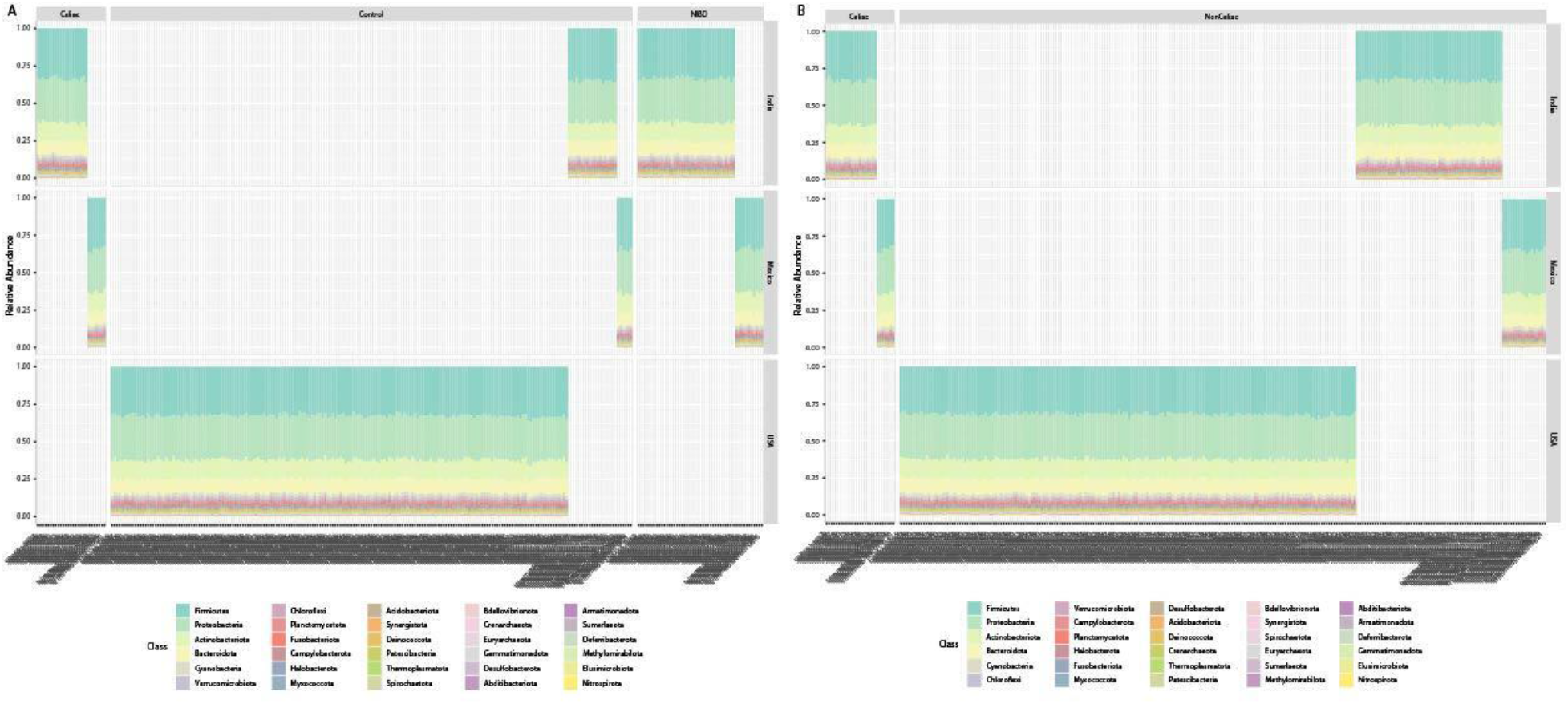
Stacked area percent abundance of normalized data. A) Control normalized percent abundance plots of control normalized data at the phylum level. B) Stacked area percent normalized abundance plots of non-CD normalized data at the phylum level. No differences in community structure were noted for either normalization technique.

### LEFSe

LEFSe identified 500 significant features across CD, NIBD and healthy samples. Healthy samples had an abundance of *Bacteroides* (2 ASVs), *Prevotella, Ruminococcaceae, Oscillospiraceae* UCG 002, *Alistipes, Lachnospiraceae* (2 ASVs), *Subdoligranulum* (2 ASVs), *Dialister*, and *Eubacterium eligens* (Figure 9A). When CD and non-CD samples were compared, 415 taxa were identified, with *Escherichia-Shigella* and *Eubacterium coprostanoligenes* being elevated in CD samples and *Lachnospiraceae* (3 ASVs), *Subdoligranulum* (2 ASVs) *Eubacterium eligens, Dialister, Alistipes, Oscillospiraceae* UCG-002, *Ruminococcaceae, Bacteroides* and *Prevotella* being elevated in non-CD samples (Figure 9B). 498 ASVs were associated with geography. *Prevotella* was elevated in Indian samples, *Oscillospiraceae* UCG-002 in Mexican samples, and *Ruminococcaceae, Alistipes* and *Bacteroides* in American samples (Figure 9C).

### Pooled Normalized Feces

Normalization of the pooled feces samples yielded 799 ASVs compared to non-normalized data, which yielded 6684 ASVs.

### Community structure analysis

Both normalization sets had lowered alpha-diversity for diseased samples using both Shannon and Simpson diversity indices (Shannon control normalized p = 0.0031876, Simpson control normalized p = 1.2784*10^−6^, Simpson non-CD normalized, Shannon non-CD normalized p = 0.020527, ANOVA Figure 10 A, B). The Simpson index of non-CD normalized data showed lowered fecal alpha-diversity for both CD and NIBD; however, these results were not significant (p = 0.12809 ANOVA). No clustering was achieved by weighted unifrac for either normalization group. Unweighted unifrac analysis plotted all samples directly on top of each other and was similarly unable to accurately cluster. To determine whether these were biologically relevant clusters, non-phylogenetic methods were utilized, with both Bray-Curtis and Jaccard indices failing to produce clusters for either normalization group (Figure 10 C, D). Stacked area percent abundance bar plots showed no difference in community structure as both a function of disease and geographic region for either normalization set (Figure 33B).

### LEFSe

LEFSe failed to identify any differentially abundant taxa between the study groups for both normalization sets.

## DISCUSSION

FDRs of CD patients are noted to have microbiomes that are atypical: distinct from the healthy microbiome and similar to the CD microbiome (for this reason both duodenal biopsies and fecal samples from Bodkhe et al. were included as NIBD samples rather than healthy, as such perturbations may confuse the analysis (15)). In any case, it was noted that dysbiotic samples (NIBD and CD) from both India and Mexico more closely resembled each other rather than healthy samples. This may be indicative of dysbiosis because of disease, rather than disease because of dysbiosis.

### Duodenal and fecal alpha- and beta-diversity analysis

Both duodenum and stool samples clustered based upon geographic region rather than disease or CD status, indicating that factors associated with geographic location are more influential than disease in terms of microbial community structure. This is not unexpected, as geographic factors heavily influence microbiome composition. Surprisingly, little differences in alpha-diversity were found between disease states (Figure 1, 7). It was previously noted that CD patients have stool and duodenal microbiomes with lowered alpha-diversity compared to healthy controls (1), but our analysis across 376 samples failed to replicate this finding. This, together with the findings from our beta-diversity/clustering, suggest that CD and healthy microbiomes do not differ as much as previously thought.

### Duodenal Community Structure Analysis

Stacked abundance bar plots showed that CD and NIBD samples, while distinct from controls, more closely resembled control groups from the same region rather than samples of the same condition from another region (Figure 2, 8). *Fusobacterium* was found to be reduced in CD patients in the filtered and scaled groups (Figure 3) but not using the normalized data. This genus of bacteria was also reduced in Mexican CD patients based on our previous reanalysis (4). It is logical for this genus to be reduced in CD patients, as it has been demonstrated to inhibit T cell responses (30) with CD being a T cell-mediated disease. Both Indian and Mexican CD samples appeared to have somewhat similar levels of *Fusobacteria* (the phylum of *Fusobacterium*). However, this phylum was absent in non-CD samples from India, indicating that the trend observed in Indian and Mexican CD samples is due to regional disease-induced dysbiosis rather than being a driver of disease. If *Fusobacterium* deficiencies indeed play a role in CD progression, it would be expected that similar levels of *Fusobacterium* would be present in both Indian and Mexican non-CD patients.

*Acinetobacter* gave the strongest signal from CD-associated microbiota (Figure 3B), perhaps uncovering an association between CD and the microbiome previously absent in Western studies. This genus has been linked to Crohn’s disease in non-Western populations (31). Previous microbiome-research found that *Acinetobacter radioresistens* positively correlated but weakly associated with high levels of C-reactive protein (32), a blood inflammatory marker that is elevated in IBD patients and is used to help distinguish from IBD (including CD) from IBS. (33, 34). Our results suggest a connection between *Acinetobacter* and the systemic inflammation observed in CD, although more research is needed to fully understand this association.

No definitive microbial signature was identified in the duodenum of CD patients from India and Mexico, however that does not mean one does not exist. The duodenum is just one of several chambers of the small intestine impacted by the disease. The duodenum has the lowest concentration of bacteria in the small intestine (35) and concentrations of bacteria increase over the length of the large intestine. Perhaps the concentration of bacteria within the duodenum may be too small to exert an effect on the host (35). Other chambers of the small intestine should also be evaluated to see whether their communities mirror the dysbiosis of the duodenum.

### Differentially abundant duodenal microbiota function: connections to dysbiosis

One pathway (P163 PWY, Figure 4) that generates both acetate and butyrate was found to be reduced in CD patients compared to controls and NIBD. Acetate is used by 95% of butyrogenic taxa and acetate concentrations directly correlate with butyrate concentrations (36). Thus, the imbalance in pathways for the production and consumption of acetate seen in CD patients may reduce butyrate production, potentially worsening inflammation and dysbiosis. While no butyrogenic taxa were identified in the pooled analysis, several were identified in the previous re-analysis (4), which is consistent with the pathway imbalance seen here.

Differences in amino acid synthesis and degradation were also observed between CD patients and controls (Figure 4). In CD patients, pathways that produce ornithine, histidine and tryptophan were decreased and pathways for glutamate catabolism were lowered. Tryptophan, glutamate, and ornithine were previously linked to CD in previous studies via analysis of peripheral sera (37). Furthermore, prior work demonstrated that tryptophan and histidine are elevated in stool samples from CD patients (38). These results are further supported by studies of dextran sulfate sodium (DSS)-induced colitis in mice, which showed an increase in several amino acids with disease, including glutamate and tryptophan (39). Data generated using PICRUSt2 serve as a proxy for what is occurring in the microbiome and what metabolites are present. This gives information as to differences in the potential to perform a given pathway but does not provide information about pathway-associated gene expression whether the metabolite is being produced or substrate consumed.

Analysis of stool samples from Indian CD patients revealed deficiencies in pathways for vitamin B12 production (14). Based on our pooled duodenum analysis, CD patients were deficient in the pathway (PWY 5188) that generates the precursor for all B12 vitamins (Figure 4). This pathway begins with glutamate and ends with tetrapyrrole, which serves as the precursors for many metal-binding compounds such as B-vitamins (cobalamin) and hemes. The presence of B vitamins results in more diverse microbial populations (40), as cobalamins and hemes are used as electron acceptors in oxygen-poor environments. Cooperative electron transport among microbes has been identified across several environments (40,41) including the gut (42). Lowered abundance of pathways needed to produce vitamin B and K, vitamin B precursors, and flavodoxin may indicate a breakdown in the shared electron transport chain in the CD microbiome. Whether this breakdown is a symptom or cause of dysbiosis cannot be determined without metabolic data and future investigation is required to fully understand the relationship between the cooperative electron transport, IBDs, and dysbiosis.

### Differentially abundant stool microbiota

Several taxa found to be less abundant in CD samples were also identified in American studies of CD stool samples (Figure 9). *Alistipes* and *Ruminococcaceae* were both less abundant in CD and NIBD stool samples and were associated with healthy samples. Another bacterium, *Escherichia-Shigella*, was identified in CD stool samples. Consistent with our findings, *Escherichia-Shigella* was previously isolated from the stools of American CD patients and shown to induce immune responses (43). *Lachnospiraceae* was also found enriched in CD stool samples (Figure 9). This bacterium was previously found to be overabundant in stool samples from children at-risk for CD (44). Our results suggest the increase in abundance of *Lachnospiraceae* persists once the disease becomes active. This is somewhat confusing, as this family of bacteria is commonly identified as a “good” bacteria. One study seeking to profile members of this family found that most members are beneficial to the host; however, some members were associated with IBD (45). This reflects a limitation of 16S rRNA-based analysis: it cannot identify some ASVs with high specificity. It is possible that there are species and strains of *Lachnospiraceae* that are associated with CD that are impossible to identify with the analysis methods we used.

### Dealing with noise and batch effects

One challenge of working with microbiome data is sheer size. ASV tables often consist of thousands of taxa, which produces noise for analysis that makes it difficult to draw conclusions. Normalization techniques have been introduced to correct for such errors. There is debate as to whether such techniques are biologically relevant, with some scientists going as far to say that results generated using any sort of transformation are not meaningful (46). In our analysis, total sum-scaling and filtering were applied and produced distinct results compared to the normalization technique used by Gibbons *et al*. Filtering and scaling identified taxa associated with disease, while normalization generated community structure data that were more in-line with the literature. Currently, it is impossible to tell which analysis technique produced the most accurate results.

### Pooled analysis and dysbiosis

Analysis of each study individually produced different results than the pooled analysis. This could be due to either noise or batch effects, both of which were corrected for and did not produce results showing that any specific disease-causing taxa were enriched across studies. Instead, we found that bacteria associated with dysbiosis or an unbalanced community structure were associated with CD. This was further supported by the results revealed by PICRUSt2 analysis, which again demonstrated connections to dysbiosis as a whole, but no strong evidence that the CD microbiome plays a causative role in disease progression. It would instead appear that the microbial community found in CD patients is a result of the disease. The procedure described in Gibbons *et al*. produced results which were distinct from these. Because the results of this method are not robust, i.e., they are not mirrored by the other techniques, this procedure may too stringently correct for batch effects. On the other hand, results from other studies do not suggest there is a CD microbial signature. While the results of the normalization described in Gibbons *et al*. are distinct from the others, they ultimately show the same thing: that the CD microbiome is not causative for disease and its differences are regionalized. Investigation and consensus are desperately needed to deduce what normalization techniques produce the most biologically relevant data.

## SUPPLEMENTARY MATERIALS AND METHODS

Studies with available data were gathered using search queries “celiac microbiome”, “celiac disease and the microbiome”, “celiac disease and gut microbiota”, and “celiac disease and gut-microbiome”. Studies which were selected examined the v4 variable region of the 16s ribosomal subunit (rRNA).

Collected sequences were prepared for dada2 (1). This was done using Cutadapt (2) and the following command “cutadapt -g ADAPTERSEQUENCE1 -g ADAPTERSEQUENCE2 -o output input”. Adapter sequences were provided by the Materials and Methods sections of the respective parent study. This was done for all studies with the exception of Bodkhe et al., in which the adapter sequences were removed using the trimLeft = c(20,20) in dada2’s filterAndTrim step. Next, the sequences were passed to dada2. We used the same steps as the original paper for Bodkhe et al., since the parent study also used dada2. Both Garcia-Mazcarro et al. and Bonder et al. used single-end sequencing; adjustments were made to the pipeline in accordance with the developer’s advice on the dada2 FAQ page for running the pipeline with single-end data. Taxonomy was assigned in dada2 using the Silva nr99 v138 training set(3).

The ASV tables from each study were then merged using dada2’s mergesequencetables. Studies with paired stool samples and biopsies were split into biopsy and stool sample sets, and analyzed separately. To create a phylogenetic tree, Clustal-o (4) was used locally, with the alignment being passed off to FastTree (5) using the -gtr and -nt options.

Next, the data generated by dada2 was prepared for PICRUSt2(6-10) (by creating an .fasta file of the ASV sequences and .biom table using the following R commands using the BIOM R package (11):

biomTable<-make_biom(t(seqtab.nochim), sample_metadata = NULL,

observation_metadata = NULL id = NULL,matrix_element_type = “int”)

write_biom(biomTable, biom_file=“table.biom”)

asvTable<-seqtab.nochim

write.table(asvTable, file=“ASVTableNewDataDADAbimera.txt”,

row.names=TRUE, sep=“\t”)

write.fasta(sequences = as.list(sequences), names = as.list(sequences), nbchar = 80, file.out = “ASV.fasta”)

This was then passed to PICRUSt2 (6-10) and run using the default parameters. The resulting data was then passed to microbiomeAnalyst (12, 13). Filtering in microbiomeAnalyst was done in accordance with each respective parent study’s methods in mind, and no transformation or refraction was performed. For weighted unifrac, unweighted unifrac, Shannon diversity index, Simpson diversity index, Chao1 diversity index, RNA seq (14-16), random forest and metagenome seq (17-18), default parameters were used. For LEFSe (19), features with a P-value (unadjusted) less than 0.1 and LDA score with an absolute value of 2 or more were identified as significant. Before filtering, ASVs without taxonomic assignment below Kingdom level were excluded. This removed 23,336 sequences. The data was filtered, removing ASVs with a count less than 4 or prevalence in less than 10% of samples. Features with a variance of less than 10% based on the interquartile range were also removed. This removed 5065 ASVs leaving 6684 ASVs for the remaining analysis. 5 samples with a library size of less than 3000 ASVs were removed, the data was then analyzed, with filtering and with total sum scaling applied, and with the procedure described in Gibbons *et al*. (20).

Analysis of pathway data was done using Bray-curtis, RNA seq, metagenomeSeq and LEFSe, all of which were run using default parameters.

A phyloseq (21) object was created and used to merge ASVs with identical taxonomy using phyloseq’s glom_taxa method. ASVs were collapsed at the genus level, leaving only ASVs with genus level assignments. This reduced the original unfiltered ASV table from 57,943 ASVs to 799. The resulting ASV table was then uploaded to microbiomeAnalyst using the same protocol as above.

## SUPPLEMENTARY RESULTS: RANDOM FOREST

### Filtered and Scaled Duodenal Random Forest

When looking at a factor of disease the model had an OOB 0.463, with class errors for CD of 0.786, 0.85 for healthy and 0.106 for NIBD. When looking at a factor of CD status the model had an error of 0.242 with class errors of 0.759 for CD and 0.0152 for non-CD. When looking at a factor of region, the model had an OOB of 0.0105 with class errors of 0 for India and 0.0182 for Mexico.

### Normalized Duodenal Random Forest

Random forest for control normalization had an OOB error of 0.66 with class errors of 0.929 for CD, 0.879 for control and 0.25 for NIBD. Random forest for non-CD normalized had an OOB of 0.309 with class error of 1 for CD and 0.0147 for non-CD. The control normalized set had an OOB of 0.0722 for geographic region with class errors of 0.143 for Indian cohort samples and 0.0182 for Mexican cohort samples. Geographic region analysis of the non-CD normalized data had an OOB of 0.0309 with class errors of 0.0476 and 0.0182 for Indian and Mexican samples respectively.

### PICRUSt 2 Duodenal Random Forest

Random forest analysis of duodenum MetaCyc pathways had an OOB error rate of 0.469 with class error rates of 0.464 for CD, 0.6 for control, and 0.394 for NIBD. Random forest using CD vs non-Cd had an OOB error rate of 0.272 with class error rates of 0.552 for CD and 0.115 for Non-CD. When looking at the region the sample was taken from, the model had an OOB error of 0.037 with class errors of 0.0769 for Indian samples and 0.0182 for Mexican samples.

### Filtered and Scaled Fecal Random Forest

Random forest had an OOB of 0.111 with class errors of 0.741 for CD, 0.0197 for healthy, and 0.143 for NIBD. For CD, versus non-CD the model had an OOB of 0.0789 with class errors of 0.815 for CD and 0 for non-CD. Random forest had an OOB of 0.208 for control normalized with class errors of 1 for CD, 0 for controls, and 0.939 NIBD. Random forest analysis of non-CD had an OOB error of 0.0968 with class error rates of 1 for CD and 0 for non-CD. When examining the region, the model had an OOB of 0.0251 with class errors of 0.0519 for Indian, 0.125 for Mexican, and 0 for American samples.

### Normalized Fecal Random Forest

Random forest analysis of control normalized data had an OOB 0.168 with class errors of 1 for CD, 0.0197 for healthy and 0.327 for NIBD. Random forest analysis of non-CD normalized data had an OOB 0.0968 with class errors of 1 for CD and 0 for non-CD.

### PICRUSt 2 Fecal Random Forest

Random forest analysis of MetaCyc pathways from fecal samples had an OOB error of

0.193 with class errors for CD being 0.963, NIBD 0.654 and control 0.0415. When looking at CD vs non-CD, the model had an OOB error 0.1 with class errors for CD being 1.0 and 0 for non-CD. When examining the geographic region, the model had an OOB error of 0.0259 with class errors of 0.0441 for Indian samples, 0.167 for Mexican samples, and 0 for American samples.

## SUPPLEMENTARY DISCUSSION

### Differentially abundant duodenal microbiota

*Fusobacterium* was found to be reduced in CD patients in the filtered and scaled (Figure 3) but not from normalized data. This genus of bacteria was also reduced in CD patients in our previous reanalysis of the Mexican data (23). It is logical for this genus to be reduced in CD patients, as it has been demonstrated to inhibit the response of T-cells (24), with CD being a T-cell mediated disease(25). Both Indian and Mexican CD samples appeared to have somewhat similar levels of *Fusobacteria* (the phylum of *Fusobacterium*). However this phylum was absent from non-CD samples from India indicating that the trend observed in Indian and Mexican CD samples is regional disease-induced dysbiosis rather than a cause of the disease. If *Fusobacterium* deficiencies indeed play a role in the progression of CD, similar levels of *Fusobacterium* would be present in both Indian and Mexican non-CD patients.

*Haemophilus* was enriched in healthy samples compared to CD samples, a confirmation of our finding in our previous work (23) (Figure 3).However, this bacteria was also found to be enriched in the Mexican samples, meaning that this genus correlated with geographic region and was lower in Indian samples. This result may be noise due to batch effects. In a previous profiling of the Indian microbiome, *Haemophilus* was not found to constitute the Indian gut microbiome in significant numbers, further supporting that this is noise(16).

Similarly, *Akkermansia* was associated with CD and region of origin, being elevated in both CD and Indian biopsies (Figure 3). This genus of bacteria was previously identified as reduced in Italian pediatric CD patients, and was noted as beneficial when found in association with the gut lining (26,27). Perhaps indicating that *Akkermansia* is not contributing to CD and that this result represents a false-positive due to a lack of healthy Indian duodenal biopsies

*Acinetobacter* gave the strongest signal from CD-associated microbiota (Figure 3B). This genus has been linked to Crohn’s disease in non-Western populations (28). Perhaps uncovering an association between CD and the microbiome absent in Western studies. Previous microbiome-research regarding post-menopausal women found that *Acinetobacter radioresistens* was positively correlated, but weakly associated with high levels of C-reactive protein. C-reactive protein is an inflammatory blood marker which has been observed as elevated in IBD patients and is used to determine whether a patient is suffering from IBD (including CD) or IBS, with IBD patients having elevated levels (29-31). This result perhaps illustrates a connection between *Acinetobacter* and the systemic inflammation observed in CD, although more research is needed to fully understand this association.

No definitive microbial signature was identified in the duodenum of CD patients from India and Mexico, however that does not mean one does not exist. The duodenum is just one of several chambers of the small intestine impacted by the disease. The duodenum has the lowest concentration of bacteria in the small intestine (32). Concentrations of bacteria increase over the length of the large intestine. Perhaps the concentration of bacteria within the duodenum may be too small to exert an effect on the host (32). Other chambers of the small intestine should also be evaluated to see whether their communities mirror the dysbiosis of the duodenum.

### Differentially abundant duodenal microbiota function: connections to dysbiosis

One pathway (P163 PWY, Figure 4) which generates both acetate and butyrate, was found to be reduced in CD patients compared to controls and NIB. A decrease in acetate production may in turn lower levels of acetate in CD patients. Acetate is used by 95% of butyergenenic taxa; and acetate concentrations directly correlate with butyrate concentrations (33). Thus, this imbalance in pathways for the production and consumption of acetate may reduce butyrate production, potentially worsening inflammation and dysbiosis. While no butyergenenic taxa were identified in the pooled analysis, several were identified in the previous re-analysis (14), this result perhaps explaining why. The fact that this pathway was found to be universally lowered in CD, despite geographic region of isolation, perhaps indicates that the metabolome of CD which should be the focus of study rather than the microbiome.

Differences in amino acid synthesis and degradation were observed with CD (34). Affected patients had elevated pathways for the production of ornithine, histidine and tryptophan, while non-CD upregulated other pathways, including ones that degrade glutamate or used glutamate as a substrate. Tryptophan, glutamate, and ornithine were previously linked to CD in previous studies; however, these results were generated using peripheral sera (34). There is little in the literature regarding whether metabolic data taken from blood sera reflects activity by the gut-metabolome; however, previous work demonstrated that tryptophan and histidine are elevated in stool of CD patients (35). Furthermore, DSS-induced colitis in murine models also showed an increase in several amino acids, including glutamate and tryptophan, further supporting the PICRUSt2 results (36). Data generated using PICRUSt2 serve as a proxy as to what is occurring in the microbiome and as to what metabolites are present. This is because PICRUSt2 aligns ASVs to reference genomes, annotates the genomes, and makes predictions on the abundance of a given pathway based on enzyme count. This gives information as to differences in the potential to perform a given pathway, but does not actually provide information as to whether those genes are being expressed and the metabolite being produced or substrate consumed. That being said, other *in-silico* studies also detected elevated amino acid synthesis in IBD-mediated dysbiosis (37).

As noted previously, stool samples from Indian CD patients were characterized by a deficiency in pathways for the production of vitamin B12 (38). In the pooled duodenum analysis, CD patients were deficient in a pathway generating the precursor to all of the B12 vitamins (Figure 4). This pathway (PWY 5188) begins with glutamate and ends with tetrapyrrole. Tetrapyrroles serve as the precursors of many metal-binding compounds, such as B-vitamins (colbamine) and hemes. As previously noted, the presence of B vitamins in a microbial community is able to induce a more diverse environment (39). Cobalamins and hemes are both used as electron acceptors in environments poor in oxygen. Cooperative electron transport among microbes has been identified across several environments (39,40) and exists within the gut (33). Lowered abundance of pathways producing vitamin B and K, vitamin B precursors, and flavodoxin may indicate a breakdown in the shared electron transport chain in the CD microbiome. Whether the breakdown is a symptom or cause of dysbiosis is impossible to know without metabolic data, so investigation into the CD metabolome should be done to verify these results.

### PICRUSt2: An improved tool

Overall, the results generated using PICRUSt2 corroborated the limited information found in the literature, showing that new computational tools are able to generate valid results from older data. More results regarding the CD metagenome and metabolome were generated using PICRUSt2 compared to PICRUSt1, which was used by many of the previous studies. This result is of no surprise given that PICRUSt2 is able to incorporate more user generated data due to its use of ASVs rather than just reference GreenGenes OTUs as is the case in PICRUSt1. Furthermore, PICRUSt2 has a reference database that is 20x larger than that of PICRUSt1, with functional information being reported as both pathways and individual KEGG modules. The pathway data generated by PICRUSt2 is regarded as the highest level output, with its predictions being made using enzyme counts (ECs). This is much more useful compared to just KEGG modules, which are the only output of PICRUSt1, as many enzymes belong to more than just a single pathway and are implicated in the generation of several metabolites. By looking at the aggregate enzyme counts, PICRUSt2 is able to better predict which pathways have the potential to be elevated in a given microbial community.

### Random forest

Random forest is a supervised method of machine learning for the classification of disease state based upon microbial abundance data. Machine learning algorithms such as random forest have been of interest; they represent a non-invasive method of diagnosis in CD. If these algorithms can accurately identify disease state from stool abundance data, with a high sensitivity, then stool sampling may represent a better diagnostic than duodenal biopsies which require patients to undergo a surgical procedure and are the gold standard for CD diagnosis. Random forest has been demonstrated to accurately predict the origin of fecal samples. For instance, in Roguet *et al*., a random forest algorithm predicted whether a given stool sample came from a cat, dog, pig, deer, or human using microbial abundance data (41). It should be noted that these communities should be expected to deviate significantly from each other as they are from entirely different host taxa. Microbiomes are highly adapted to their host, and it should be of no surprise that these communities are easy to tell apart. Our implementation of random forest analysis was able to accurately predict the country of origin of both the duodenal and fecal pooled analysis, however across the studies and pooled analysis, it was unable to accurately predict disease status. It is known that geographic region can have drastic impacts on the composition of microbial communities within the host due to a variety of factors including diet (42, 43). This may illustrate that despite having different disease states, CD and NIBD patients have a similar microbiome to healthy individuals of the same population as opposed to CD or NIBD patients from other continents. This is further supported by unifrac analysis of the individual studies and Bray-Curtis analysis of the pooled studies which illustrated that samples tended to cluster together on the basis of country of origin rather than disease. This does not however mean that random forest classification is not applicable to IBD and CD, with Chehoud *et al*. being able to accurately predict IBD status using both bacterial and fungal data, perhaps demonstrating the importance of fungi within the gut microbiome. If anything, this result reinforces the idea that the differences observed in CD community structure are not uniform and likely do not play a contributing role in the progression of the disease.

### Previous attempts unify CD knowledge

There are several sources of information describing gut-dysbiosis associated with celiac disease, however these resources are limited either in quantity of information, or to a single region of the world. One such study (44) examined 500 infants at risk for the development of celiac disease over the course of five years to identify changes in the gut-microbiota and metabolome associated with the development of celiac disease. While this study did include a large sample size, it focused on developed Western countries, and is representative of those populations. This does not give an accurate representation of all the potential bacteria which could be associated with CD as diet, antibiotic use and other factors associated with Western countries have a large impact on microbiome composition. Furthermore, this study utilized mostly stool and blood samples, which serve as proxies for studying CD, which is mainly active in the duodenum. There are also existing databases which serve to unify data from other studies under one archive, gutMDisorder is one such resource. GutMDisorder is a good start, but is lacking in information on CD with only eleven entries (under categories “celiac disease”-7 entries, “coeliac disease”-1 entry and “gluten-free diet”-3 entries) as relating to the CD microbiome, of which only three had information regarding the species level (45). This is problematic as associations above the genus level are rather uninformative as the function of bacteria can vary wildly even at levels as low as strain; thus information about genus and higher classification, while useful in exploration of ecosystem structure, inform little regarding mechanisms of disease development or targets of intervention. Thus more work is needed to unify information taken from previous studies under a single resource to better understand the relationships between CD and the microbiome.

### Dealing with noise and batch effects

One challenge of working with microbiome data is its sheer size. ASV tables often consist of thousands of taxa, producing considerable noise for analysis and making drawing conclusions from such data difficult. Normalization techniques have been introduced to correct for such errors. There is debate as to whether such techniques are biologically relevant with some scientists going as far to say that results generated using any sort of transformation are not meaningful (46). In our analysis, total sum-scaling and filtering were applied producing distinct results compared to the normalization technique taken from Gibbons *et al*. Filtering and scaling did identify taxa associated with disease, however normalization produced data regarding community structure which was more in-line with the literature. Currently, it is impossible to tell which analysis technique produced the most accurate results.

### Pooled analysis and dysbiosis

Each study individually produced different results than the pooled analysis. This could be due to either noise or batch effects, both of which were corrected for and did not produce results showing that any specific disease causing taxa was enriched across studies, but instead showed that bacteria associated with dysbiosis, or an unbalanced community structure were associated with CD. This was further amplified by the results of the PICRUSt2 analysis, which again demonstrated connections to dysbiosis as a whole, but no real evidence that the CD microbiome plays a causative role in the progression of the disease. It would instead appear that the microbial community found in CD patients is a result of the disease. The procedure described in Gibbons *et al*. produced results which were distinct from these. As the results of this method are not robust, *i*.*e*. they are not mirrored by the other techniques, this procedure may be too stringent in its attempts to correct for batch effects. On the other hand, the results of the other studies do not seem to indicate that there is a CD microbial signature. Perhaps indicating that while the results of the normalization described in Gibbons *et al*. are distinct from the others, they are ultimately showing the same thing, that the CD microbiome is not causative of the disease and its differences are regionalized. Investigation and consensus is desperately needed to deduce what normalization techniques produce the most biologically relevant data.

### Pooled meta-analyses generate different results from parent studies

A previous meta-analysis (47) of sequencing data from colorectal cancer stool and tissue samples also found that features associated with disease were not uniform across samples, but rather had a patchy distribution, with some studies having a strong signal indicating that a taxa was highly associated with the disease, while in other studies the signal was reduced or absent. To these researchers, this indicated that these taxa may be associated with the disease, and while some may worsen patient outcomes, are not a required component of the mechanism of pathogenesis in colorectal cancer. While this study was conducted on an entirely separate disease, the same may be true of CD, with some “bad” taxa associating with the disease and worsening symptoms and recovery but not actually playing a casual role in the prognosis of a patient from inactive to active CD.

## CONCLUSIONS

### Regional differences in CD

Our results show that, while the CD microbiome is indeed distinct from that of the healthy microbiome, diseased samples are more similar to those taken from the same region rather than those from CD patients from a different region, which argues that dysbiosis seen in CD patients is a result of the disease rather than a driver of it. Thus, one difficulty in interpreting our data was that people from different parts of the globe had location-specific microbial profiles, making it hard to compare CD cohorts from different regions. Our previous analysis (14) showed that certain taxa are enriched or deficient in CD, but our pooled analysis was not able to detect these changes, again indicating these differences are likely a result of the disease rather than a contributing factor.

### Metabolic differences in CD

Our previous analysis found that pathways needed to produce electron-accepting products were reduced in the CD microbiome (4). Here we found that precursors to these pathways are reduced across geographic boundaries. We also found perturbations in amino acid synthesis in CD patients, which was also confirmed by other CD microbiome studies. These results confirm that analysis using PICRUSt2 generates valid predictors of microbial function.

### Small intestinal biopsies versus stool samples

Our studies focused on biopsies taken from the duodenum and stool samples. CD is active in the small intestine, meaning that stool samples are likely not reflective of the chambers of the gut where the CD is active. The duodenum represents a more logical choice of study for the CD microbiome, but it is not the only chamber of the small intestine affected. While we found no microbial signature associated with CD duodenal samples in this analysis, that does not mean that the small intestinal microbiome is not involved. Characterization of the jejunal and ileal microbiome should also be pursued.

### Summary

Overall, our findings indicate that the dysbiosis observed in CD is likely a result of the disease rather than a contributing factor. Analysis of data from any geographic region individually produces results showing potentially relevant differentially abundant taxa. However, these results do not hold up across pooled analysis, suggesting they are not contributing to the disease or contribute to the disease in a specific cohort in a manner that is not generalizable to the global population. This is supported by the PICRUSt2 functional data, with connections to other traits observed in dysbiotic communities.

## SUPPLEMENTARY FIGURES

### SUPPLEMENTARY FIGURE LEGENDS

**Supplementary figure 1.**
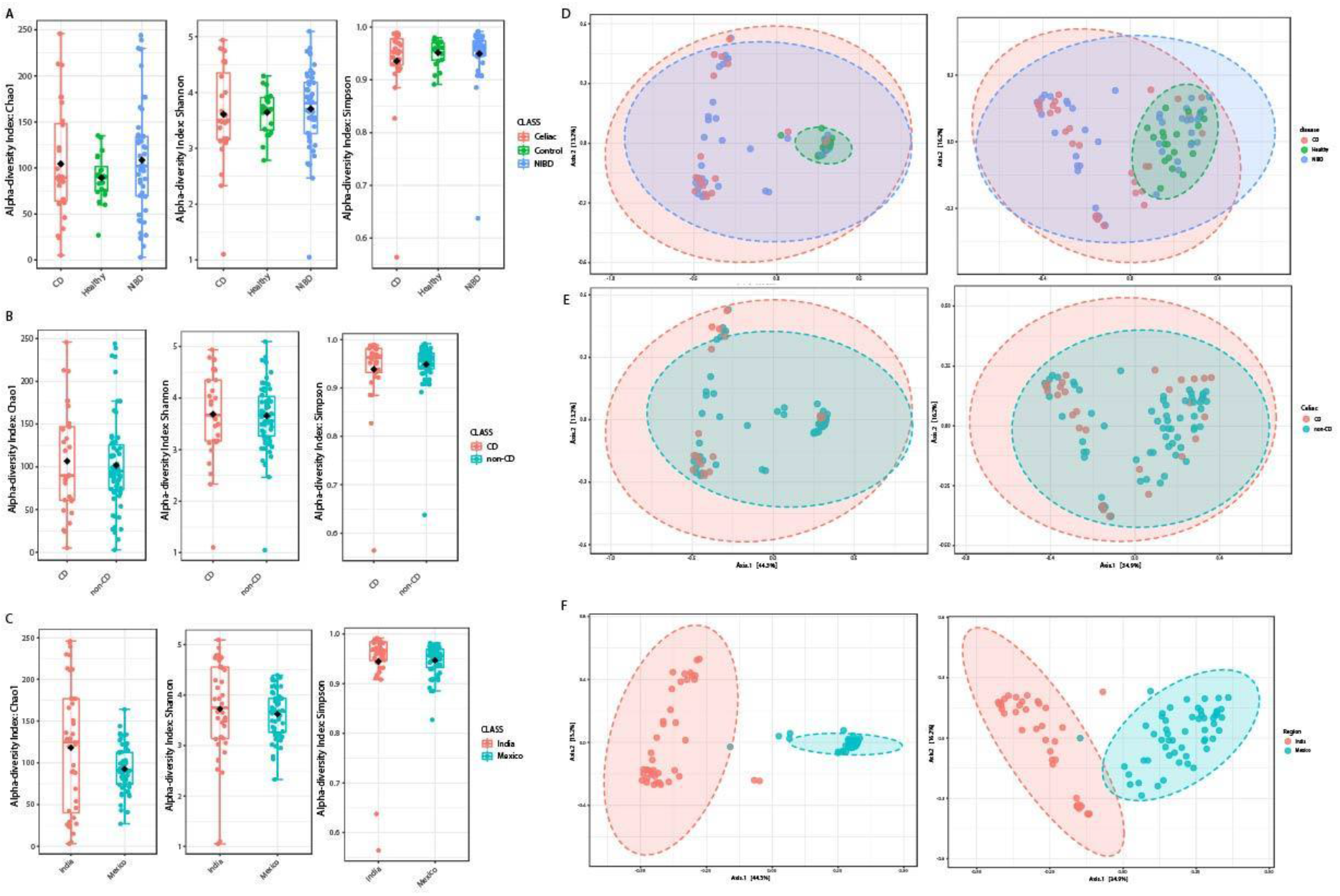
pooled filtered and scaled duodenal alpha and beta diversity analysis: A) Alpha-diversity as a factor of disease, with red representing CD, green healthy and blue NIBD. No significant difference was noted for Chao1, Shannon or Simpson (P >0.05) .B Alpha-diversity as a factor of CD status with red representing CD and blue non-CD. No significant difference was noted (P >0.05). C) Alpha-diversity as a factor of region, with red representing Indian samples and blue representing Mexican samples. Indian samples were significantly enriched for Chao1 (P = 0.027265) but not Shannon or Simpson (P > 0.05). D) Unweighted (left) and weighted (right) unifrac analysis of CD (red) NIBD(blue) and Healthy(green). E) Unweighted (left) and weighted (right) unifrac analysis of CD(red) versus non-CD (blue). F) Unweighted (left) and weighted (right) unifrac analysis, with Indian samples in red and Mexican in blue. All unifrac analysis yielded P-values less than 0.001.

**Supplementary figure 2:**
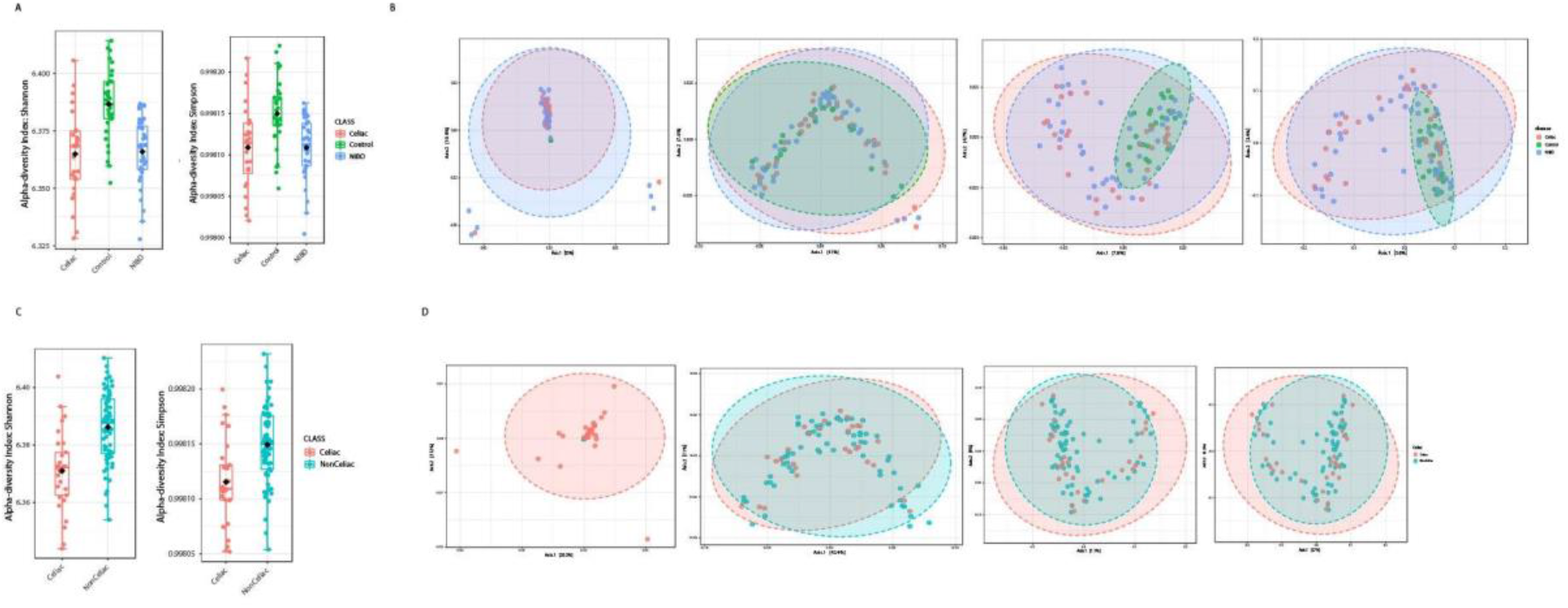
Normalized duodenum alpha and beta diversity. A) Control normalized alpha diversity. The left plot shows group Shannon diversity for the control normalized dataset with disease samples having lower alpha-diversity compared to healthy samples. This was observed in both Shannon (right) and Simpson (left) indices (hannon control normalized P = 7.8156*10-8, Simpson control normalized P = 7.8156*10-5). Red shows CD patients, green healthy and blue NIBD. B) shows clustering methods, with the left most being unweighted unifrac, followed by weighted unifrac, Bray-Curtis, and Jaccard index. No significant clustering was achieved on the basis of disease for any of the methods (P> 0.05). C) shows non-CD normalized alpha diversity with Shannon index on the left and Simpson on the right, with red showing CD patients and blue non-CD patients. CD patients had significantly lower alpha-diversity compared to non-CD patients for both tests (Simpson non-CD normalized P= 0.0001319, Shannon non-CD normalized P = 0.034427). D) Beta-diversity of non-CD normalized data, with unweighted unifrac on the far left, followed by weighted unifrac, Bray-Curtis and Jaccard indices. No significant clustering was achieved on the basis of CD status for any method (P >0.05).

**Supplementary figure 3.**
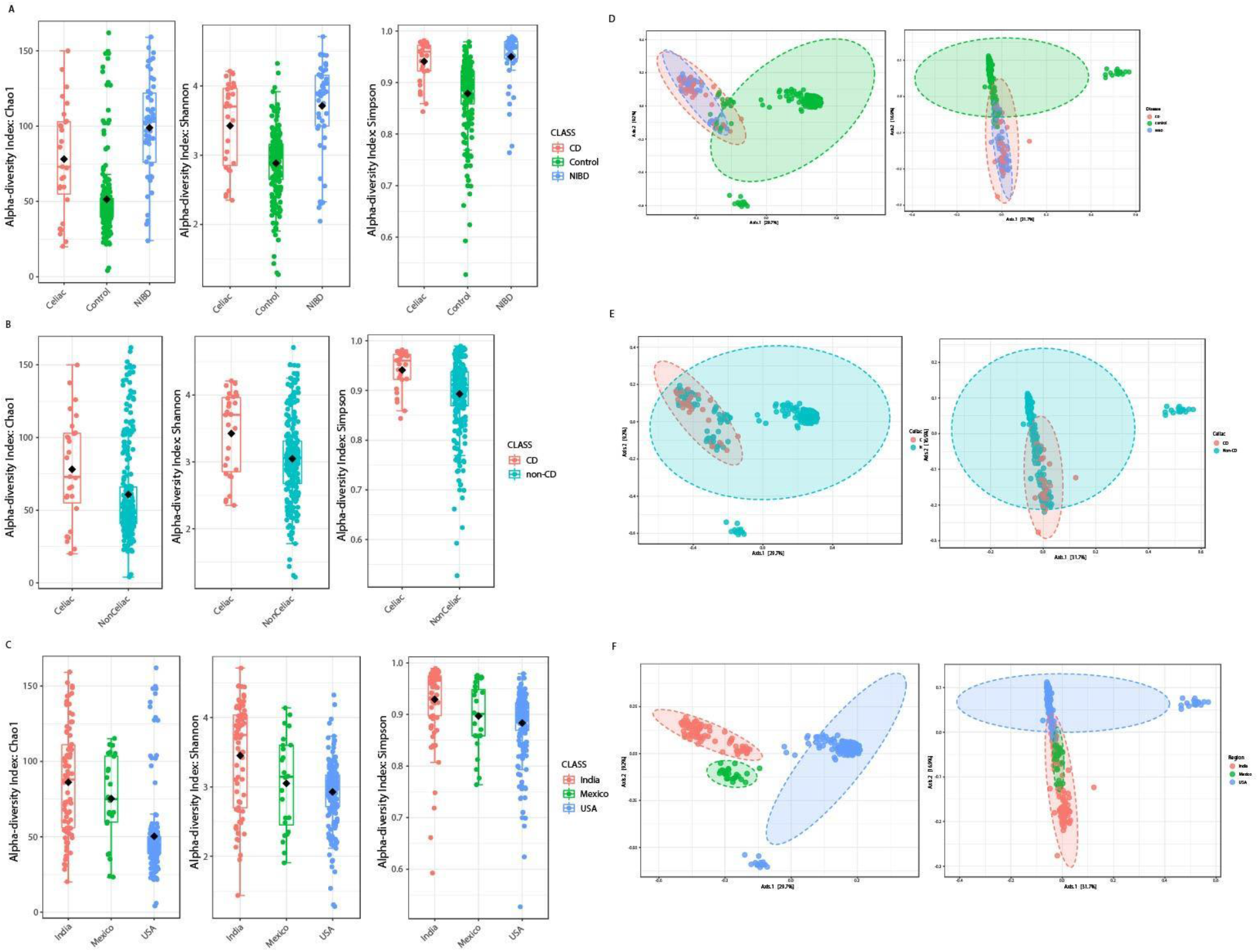
Filtered and scaled fecal alpha and beta diversity analysis. A) Alpha diversity of CD (red), healthy (green) and NIBD (Blue) stool samples. NIBD and CD were found to have higher alpha-diversity based on the Chao1(left), Shannon (middle) and Simpson (right) indices (P = 7.1026*10-28, P = 8.0086*10-23, P = 3.3789*10-13). B) Alpha diversity of CD versus non-CD showed that CD (red) had higher alpha diversity than non-CD (blue) across the Chao1 (right), Shannon (middle) and Simpson (right) indices (P =0.0090813, P = 0.0015761, P = 3.357*10-7). C) Alpha diversity as a function of region showed that American samples (blue) had the lowest alpha diversity compared to Mexican (green)and Indian(blue) samples (P = 3.1866*10-21, P= 1.31478*10-11, P= 8.9231*10-7). D) shows unweighted (left) and weighted (right) unifrac analysis of CD (red), healthy (green) and NIBD (blue). No clustering as a function of disease was observed. E) Unweighted (left) and weighted (right) unifrac analysis of CD (red) versus non-CD (blue). No significant clustering was achieved as a result of CD. E) unweighted (left) and weighted (right) unifrac analysis of Indian (red), American (blue), and Mexican (green) stool samples. Clustering was achieved as a result of region for unweighted unifrac (P <0.001), but not weighted unifrac analysis.

**Supplementary figure 4.**
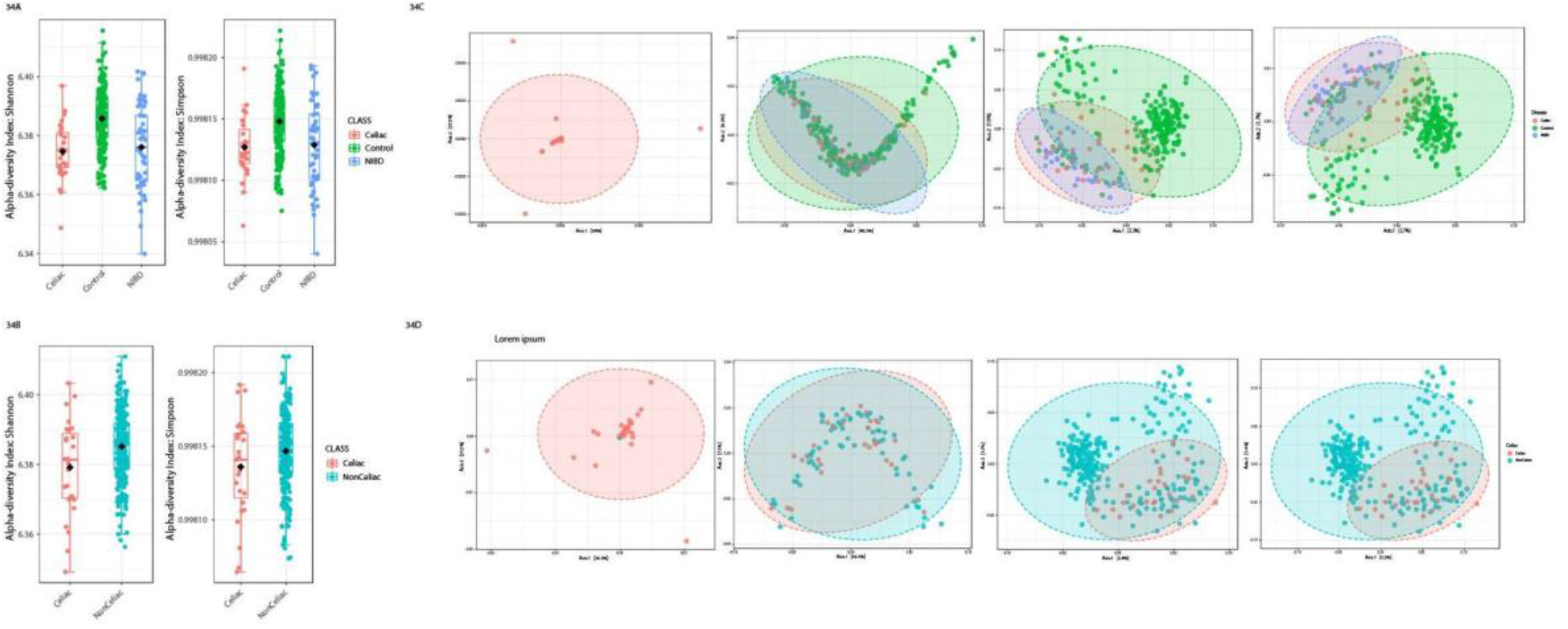
Normalized fecal alpha and beta diversity: A) Disease alpha diversity of control normalized samples. Shannon (left) and Simpson (right) alpha diversity of CD (red), healthy (green) and NIBD (blue). Healthy stool samples had higher alpha diversity compared to diseased samples (Shannon control normalized p = 0.0031876, Simpson control normalized P = 1.2784*10-6). B) CD versus non-CD alpha diversity of non-CD normalized samples. Shannon (left) and Simpson (right) alpha-diversity of CD (red) versus non-CD samples (blue). CD samples had lower alpha-diversity compared to non-CD, though this was only significant for the Shannon index (non-CD normalized P = 0.020527) C) Beta-diversity analysis of control normalized samples with unweighted unifrac (far left) followed by weighted unifrac, Bray-curtis and Jaccard indices. No significant clustering as a function of disease was identified. D) Beta-diversity analysis of non-CD normalized samples with unweighted unifrac (far left), weighted unifrac, Bray-Curtis and Jaccard indices. No significant clustering was achieved with any of the analysis methods.

